# PBP1A directly interacts with the divisome complex to promote septal peptidoglycan synthesis in *Acinetobacter baumannii*

**DOI:** 10.1101/2022.09.26.509500

**Authors:** Katie N. Kang, Joseph M. Boll

## Abstract

The class A penicillin-binding proteins (aPBPs), PBP1A and PBP1B, are major peptidoglycan synthases that synthesize more than half of the peptidoglycan per generation in *Escherichia coli*. Whereas aPBPs have distinct roles in peptidoglycan biosynthesis during growth (i.e., elongation and division), they are semi-redundant; disruption of either is rescued by the other to maintain envelope homeostasis and promote proper growth. *Acinetobacter baumannii* is a nosocomial pathogen that demonstrated a high propensity to overcome antimicrobial treatment. *A. baumannii* encodes both PBP1A and PBP1B (encoded by *mrcA* and *mrcB*, respectively), but only *mrcA* deletion decreased fitness and contributed to colistin resistance through inactivation of lipooligosaccharide biosynthesis, indicating that PBP1B was not functionally redundant with PBP1A activity. While previous studies suggested a distinct role for PBP1A in division, it was unknown if its role in septal peptidoglycan biosynthesis was direct. Here, we show that *A. baumannii* PBP1A has a direct role in division through interactions with divisome components. PBP1A localizes to septal sites during growth, where it interacts with the transpeptidase, PBP3, an essential division component that regulates daughter cell formation. PBP3 overexpression was sufficient to rescue the division defect in Δ*mrcA A. baumannii*; however, PBP1A overexpression was not sufficient to rescue the septal defect when PBP3 was inhibited, suggesting their activity is not redundant. Overexpression of a major DD-carboxypeptidase, PBP5, also restored the canonical *A. baumannii* coccobacilli morphology in Δ*mrcA* cells. Together, these data support a direct role for PBP1A in *A. baumannii* division and highlights its role as a septal peptidoglycan synthase.

**Importance:** Peptidoglycan biosynthesis is a validated target of β-lactam antibiotics, and it is critical that we understand essential processes in multidrug resistant pathogens such as *Acinetobacter baumannii*. While model systems such as *Escherichia coli* have shown that PBP1A is associated with side wall peptidoglycan synthesis, we show herein that *A. baumannii* PBP1A directly interacts with the divisome component PBP3 to promote division, suggesting a unique role for the enzyme in the highly drug resistant nosocomial pathogen. *A. baumannii* demonstrated unanticipated resistance and tolerance to envelope-targeting antibiotics, which may be driven by rewired peptidoglycan machinery, and may underlie therapeutic failure during antibiotic treatment.

## Introduction

The Gram-negative cell envelope consists of an inner and outer membrane lipid bilayers, separated by a periplasmic space enriched with peptidoglycan. The peptidoglycan sacculus determines bacterial cell shape. Peptidoglycan biosynthesis and assembly is coordinated by the elongasome and divisome, two multiprotein complexes that regulate rod length and daughter cell formation, respectively. Dogma suggested that elongation and division activities are dependent on lipid II substrate availability (1–4); therefore, rod shape and septum assembly is dictated by substrate competition between the two peptidoglycan synthase complexes. Increased elongasome activity favors a narrow, elongated rods (5) with V-shaped division constrictions (4), whereas increased divisome activity results in wide, short cells (5) with blunted septal sites (4).

In Proteobacteria like *Escherichia coli*, the divisome and elongasome complexes include more than 20 proteins that tightly regulate peptidoglycan biogenesis (6, 7). They include several regulatory components, the *<u>s</u>*hape, *<u>e</u>*longation, *<u>d</u>*ivision and *<u>s</u>*porulation (SEDS)-family glycosyltransferases, FtsW and RodA, and class B penicillin-binding protein transpeptidases, PBP2 and PBP3 that synthesize peptidoglycan along the cell axis and septum, respectively (8– 12). Class *<u>A p</u>*enicillin-*<u>b</u>*inding *<u>p</u>*roteins (aPBPs), PBP1A and PBP1B (encoded by *mrcA* and *mrcB*, respectively) are primary peptidoglycan synthases that also contribute to side wall and septal peptidoglycan biosynthesis. aPBPs are bifunctional, with distinct domains that catalyze either transpeptidase or glycosyltransferase activities. In *E. coli*, PBP1A and PBP1B are functionally semi-redundant. Individual *mrcA* and *mrcB* deletions do not contribute to measurable morphological defects (13, 14) and only one is required for viability (15). Importantly, only one aPBP is required for elongation (10) and division (16) in the well-studied model organism.

aPBPs directly interact with specific monofunctional transpeptidases in the elongasome or divisome. In *E. coli*, PBP1A associates with PBP2 in the elongasome (14). PBP2 also interacts with the monofunctional elongation glycosyltransferase, RodA (17). Elongasome activity is regulated by MreBCD through interactions with the PBP2 cytoplasmic domain (17). In contrast, PBP1B associates with divisome transpeptidase, PBP3 (18, 19), and with FtsN (18), an essential bitopic membrane protein necessary to promote division (20) by inducing PBP3 activity (21). PBP1B forms a trimeric complex with PBP3 and FtsW, the monofunctional divisome glycosyltransferase homolog of RodA (19). FtsW inhibits PBP1B-mediated peptidoglycan polymerization in the absence of PBP3 (19). While PBP1A and PBP1B interact with distinct peptidoglycan assembly complexes, either aPBP can compensate for the other to restore missing activity and function (14).

Our understanding of peptidoglycan synthases in Gram-negative bacteria are largely based on studies in the rod-shaped model organism, *E. coli*. However, accumulating evidence clearly shows that PBP1B activity is not functionally redundant with PBP1A in the highly drug resistant nosocomial pathogen, *A. baumannii* (22–24). Deletion of *mrcA*, which encodes PBP1A, caused septation defects, which induced cell chaining and cell filamentation. Moreover, PBP1A catalytic activity was necessary for completing septation (23). Consistent with a role in division, PBP1A was enriched at the midcell during growth, where divisome components assemble to regulate cell envelope invagination during cytokinesis. While phenotypes were consistent with a role for PBP1A in *A. baumannii* division, it was not determined if PBP1A directly or indirectly interacted with the divisome machinery. Here, we show that *A. baumannii* PBP1B is not functionally redundant with PBP1A. Instead, PBP1A showed distinct localization at the midcell prior to division, where it presumably promotes septal peptidoglycan synthase activity. PBP1A complexes with PBP3 during growth, indicating it directly contributes to divisome activity and daughter cell formation in *A. baumannii*. In contrast, we were unable to confirm direct interaction between PBP1A and PBP2 during growth. Further supporting an overlapping role between PBP1A and PBP3 enzymatic activity in *A. baumannii* PBP3 overexpression rescued the Δ*mrcA* division defect. In contrast, PBP1A could not rescue the septal defect when PBP3 was inhibited with aztreonam, suggesting that PBP1A and PBP3 have distinct roles in septal peptidoglycan biosynthesis. Together, these studies uncover a unique role for PBP1A in *A. baumannii* cell division and fitness.

## Results

### PBP1B is not functionally redundant with PBP1A in *A. baumannii*

In contrast to the proposed aPBP peptidoglycan synthase model in rod-shaped *E. coli*, data suggested that PBP1A has a primary role in division in coccobacilli-shaped *A. baumannii*, and PBP1B does not compensate when it is inactive (23). *mrcA* mutation induced an *A. baumannii* growth defect, cell chaining, and reduced fitness, whereas the Δ*mrcB* mutation did not. Consistent with these data, Δ*mrcA A. baumannii* expressing a transpeptidase-defective PBP1A_S459A_ (but not Δ*mrcB* or Δ*mrcB* expressing PBP1B_S455A_) impaired growth (24). To determine if *A. baumannii* PBP1B could rescue the Δ*mrcA* septation defect, we visualized cells using phase and fluorescence microscopy (**Fig 1A**). As previously done (23, 25–27), cells were grown to logarithmic growth phase and stained with the fluorescent D-alanine derivative, NADA, to visualize peptidoglycan. NADA is incorporated into peptidoglycan by PBPs and LD-transpeptidases (28–31). Consistent with previous reports (23), Δ*mrcA* cells produced multiseptated chained cells relative to wild type, while Δ*mrcB* cells did not. Using an IPTG-inducible PBP1B construct (**Fig S1A**), denoted as pPBP1B_OE_, overexpression was not sufficient to rescue the septation defect in Δ*mrcA* cells (**Fig 1A**). Length quantifications also showed that PBP1B overexpression could not restore the septation defect in Δ*mrcA A. baumannii* (**Fig 1B**).

**Figure 1.**
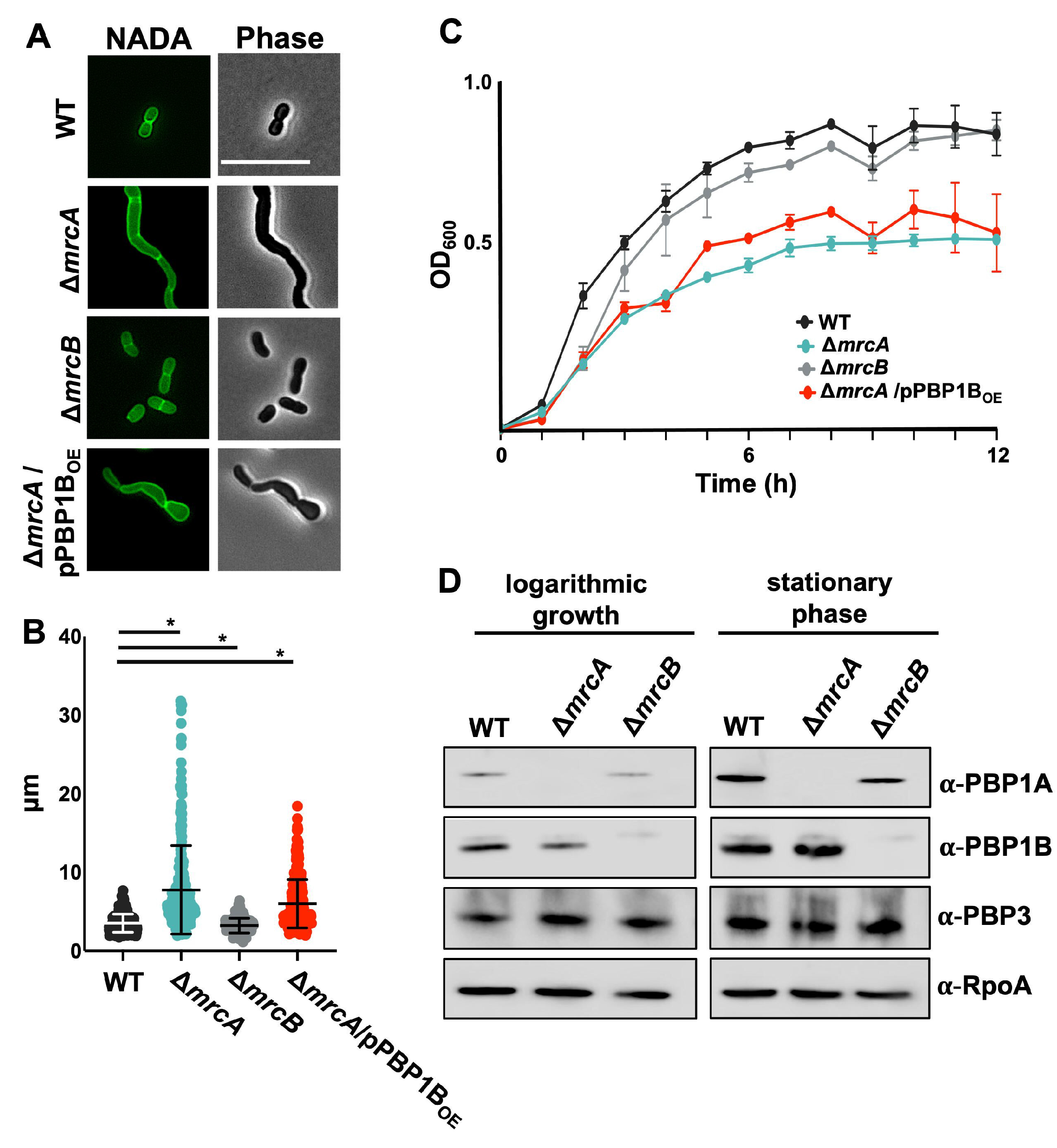
PBP1B cannot compensate for the division defect in Δ*mrcA A. baumannii*. (A) Fluorescence (left) and phase (right) microscopy of wild type (WT), Δ*mrcA*, Δ*mrcB* and Δ*mrcA*/PBP1B_OE_. Scale bar 10 μm. (B) Length (pole to pole) quantifications of each cell population (*n* ≥ 300) was calculated using ImageJ software with MicrobeJ plugin. Each dot represents one cell. Error bars represent standard deviation. Significance testing conducted using Student *t* test with two-tailed distribution assuming equal variance. *P* < 0.05. (C) Optical density growth curve of WT, Δ*mrcA*, Δ*mrcB* and Δ*mrcA*/PBP1B_OE_. Error bars represent standard deviation. (D) Western blot of WT, Δ*mrcA* and Δ*mrcB* whole cell lysates collected in mid-logarithmic growth (left) and stationary phase (right). PBP1A is 94.74 kDa; PBP1B is 88.21 kDa; PBP3 is 67.66 kDa; RpoA is 37.62 kDa.

To determine if PBP1B expression could rescue the fitness defect in Δ*mrcA* cells, growth rates of aPBP mutants were measured (**Fig 1C**). Δ*mrcB* cells did not show significant growth defects relative to wild type, and PBP1B expression was unable to rescue the growth defect in Δ*mrcA A baumannii*.

Lastly, if PBP1B could compensate for loss of PBP1A, we hypothesized that increased PBP1B levels might be evident in Δ*mrcA* cells. Using specific antisera, relative PBP1A and PBP1B levels were compared in wild type, Δ*mrcA*, and Δ*mrcB* cultures in logarithmic growth or stationary phase (**Fig 1D**) and quantified (**Fig S1B**). There was an obvious difference in aPBP levels between growth phases, where PBP1A and PBP1B expression increased in stationary phase relative to growth. Notably, PBP1A levels were reduced in Δ*mrcB* cells relative to wild type. Conversely, PBP1B levels were decreased in Δ*mrcA* cells. The changes in protein levels were reproducible and might illustrate a mechanism to maintain proper stoichiometry between the two proteins, particularly if their properties are antagonistic. While it is possible that aPBP enzyme activity could increase when the other is defective, these data support a model where the PBP1B activity is not functionally redundant with PBP1A; specifically, PBP1B cannot restore Δ*mrcA* septation and fitness defects.

### Increased PBP1A activity shifts peptidoglycan biosynthesis towards division in *A. baumannii*

Using a previously described (23) pPBP1A-mCherry reporter fusion, expressed from its native promoter (**Fig S2A**) that fully complemented the division defect in Δ*mrcA A. baumannii*, we characterized PBP1A localization. Expression showed diffuse PBP1A localization throughout the cell, but fluorescence intensity was enriched at the midcell, where divisome proteins assemble to regulate septal formation and division (**Fig 2A**). PBP1A-mCherry localization relative to septal peptidoglycan formation was compared using demographs (**Fig 2B**). PBP1A-mCherry accumulated at the midcell prior to septal peptidoglycan assembly, suggesting that PBP1A not only contributes to septal peptidoglycan biogenesis, but it could also shift peptidoglycan synthesis from elongation to division.

**Figure 2.**
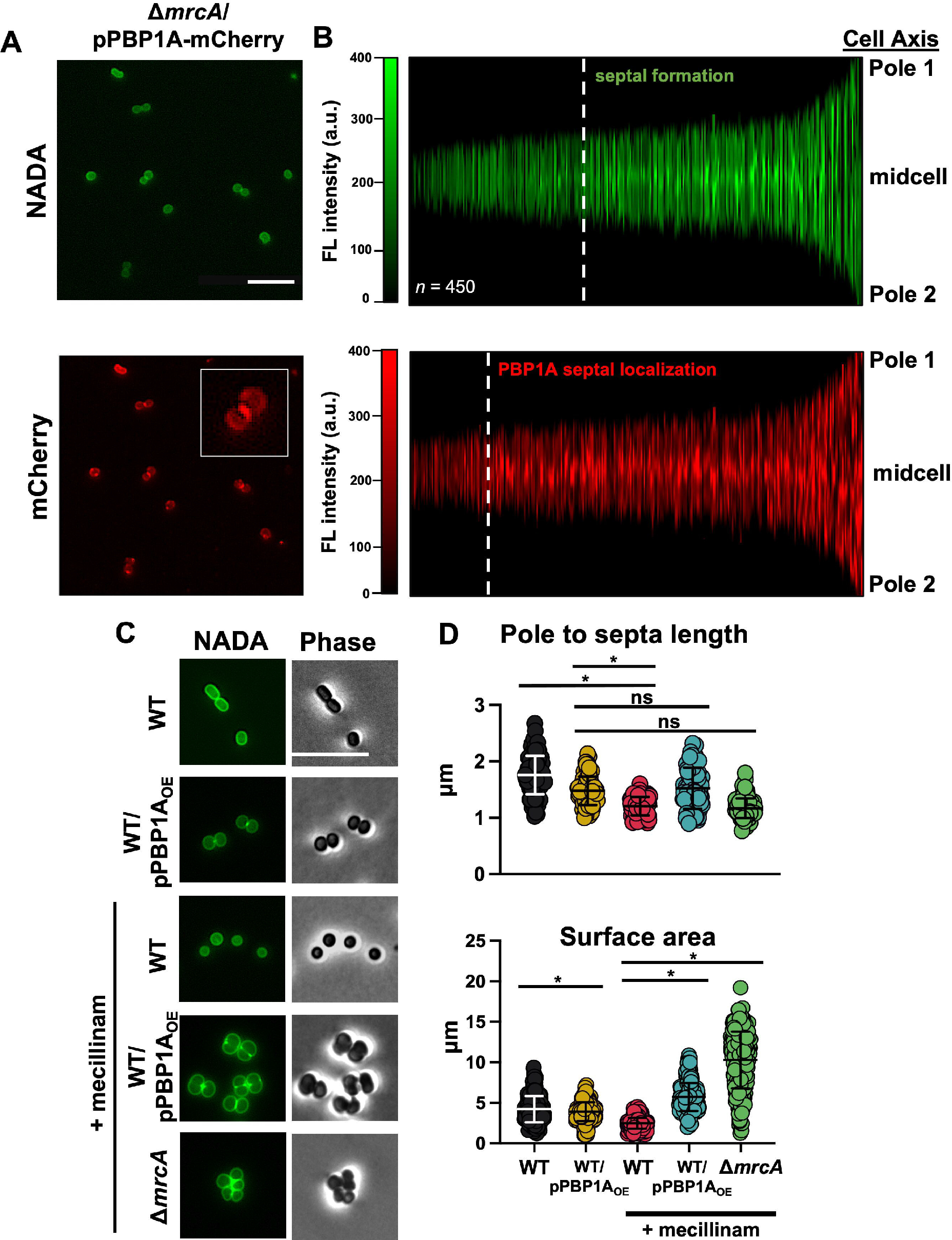
PBP1A shifts the peptidoglycan synthesis balance towards division. (A) Fluorescence microscopy of Δ*mrcA/*pPBP1A-mCherry cells, NADA fluorescence (top) and PBP1A-mCherry (bottom). PBP1A-mCherry is expressed from its native promoter. Scale bar = 10μm. The inset shows a representative cell at higher magnification. (B) Demographs depicting NADA fluorescence (top) and PBP1A-mCherry (bottom) intensity localization along the cell axis. Cells are ordered by increasing length (*n* = 450). Dashed lines indicate when midcell localization of NADA and PBP1A-mcherry becomes apparent. (C) Fluorescence and phase microscopy of wild type (WT), WT/PBP1A_OE_ (overexpression; inducible promoter), WT, WT/PBP1A_OE_, and Δ*mrcA* treated with 0.5-MIC mecillinam. Scale bar = 10 μm. (D) Length (pole to septa in dividing cells or pole to pole in nondividing cells) (*n* = 100) was calculated using ImageJ software. Area was calculated as surface square pixels in MicrobeJ. Each dot represents one cell. Error bars indicate standard deviation. Significance testing conducted using Student *t* test with two-tailed distribution assuming equal variance. * *P* < 0.05; ns is not significant.

In line with previous work (23), these data also show that PBP1A overexpression (via pPBP1A_OE_) induced cell rounding, a phenotype consistent with an increased septation rate and/or decreased elongation (**Fig 2C**). Notably, PBP2 inhibition also produced rounded cells in Enterobacterales (32). Based on these data, we hypothesized that PBP1A activity promotes septation and/or antagonizes elongasome activity in *A. baumannii*.

PBP2 is specifically inhibited by the β-lactam, mecillinam (32), so *A. baumannii* treated with sub-minimum inhibitory concentrations of mecillinam expressing either native levels or overexpressing PBP1A (**Fig S2B**), which also restored growth (**Fig S2C)**, were analyzed using fluorescence and phase microscopy (**Fig 2C**). The distance from pole to septa (or pole to pole in nondividing cells) was measured among populations (*n* = 100) to quantitate (**Fig 2D**). Intriguingly, wild type mecillinam-dependent PBP2 inhibition produced statistically significant shorter cells relative to wild type cells overexpressing PBP1A (**Fig 2D, top**). While this could be a dosage effect, where PBP2 inhibits the elongasome more potently than PBP1A, it was not clear if PBP1A directly inhibited elongasome activity. Notably, cells treated with mecillinam and overexpressing PBP1A were larger relative to mecillinam-treated wild type cells and cells overexpressing PBP1A (**Fig 2D, bottom**). While an increase in cell surface area could suggest a synthetic effect from PBP1A overexpression in combination with mecillinam treatment, an increased rate of septal peptidoglycan polymerization could also account for the increase in cell size, where cells are rapidly dividing before the prior division cycle is complete. To distinguish if cells overexpressing PBP1A produced more cells overall, we calculated colony forming units (CFU/ml) over time (**Fig S2D**). Cells overexpressing PBP1A reproducibly showed increased CFU/ml relative to wild type regardless of mecillinam treatment, suggesting that increased septation may drive formation of short round cells. In line with these data, Δ*mrcA* cells treated with sub-MIC mecillinam were round and clumped (**Fig 2C**), supporting a model where defects in both septation and elongation reduced CFU relative to wild type (**Fig S2D**). Together, these data suggest PBP2 inhibition was not sufficient to shift peptidoglycan biosynthesis towards division without PBP1A-dependent septal peptidoglycan activity.

### PBP1A directly interacts with PBP3 at the divisome during growth

Previous work (23) and the PBP1A localization studies here, are consistent with the accumulation of unproductive septal events in Δ*mrcA* cells, which implies that PBP1A promotes proper *A. baumannii* division. However, it was unclear if PBP1A directly interacted with divisome proteins to promote septal peptidoglycan biosynthesis. Next, we tested if PBP1A directly interacted with PBP2 and PBP3 *in vivo* using *<u>co</u>*-*i*mmuno*<u>p</u>*recipitation (CoIP) (33). A C-terminal Flag fusion (PBP1A-FLAG) was expressed (**Fig S3A**) in wild type. Cells were incubated with Lomant’s reagent (dithiobis succinimidylpropionate; DSP). DSP is a membrane-permeable crosslinker that reacts with primary amine groups and contains a 12.0 Å spacer arm. After DSP crosslinking, PBP1A immunoprecipitation showed direct interaction with PBP3, but not PBP2, in *A. baumannii* during growth (**Fig 3**). Furthermore, immunoprecipitation of PBP3-FLAG expressed in wild type cells (**Fig S3A**) also showed direct interaction with PBP1A (**Fig 3**). PBP3 is a highly conserved divisome protein, known to contribute to daughter cell formation during septation (34, 35). Direct interaction between PBP1A and PBP3 during growth supports a model where PBP1A directly promotes *A. baumannii* septation. Contrary to rod-shaped bacteria, where PBP1A interacts with PBP2 to promote elongasome-dependent side wall peptidoglycan biosynthesis, our data strongly suggest that PBP1A directly interacts with the divisome to promote septal peptidoglycan biosynthesis in *A. baumannii*.

**Figure 3.**
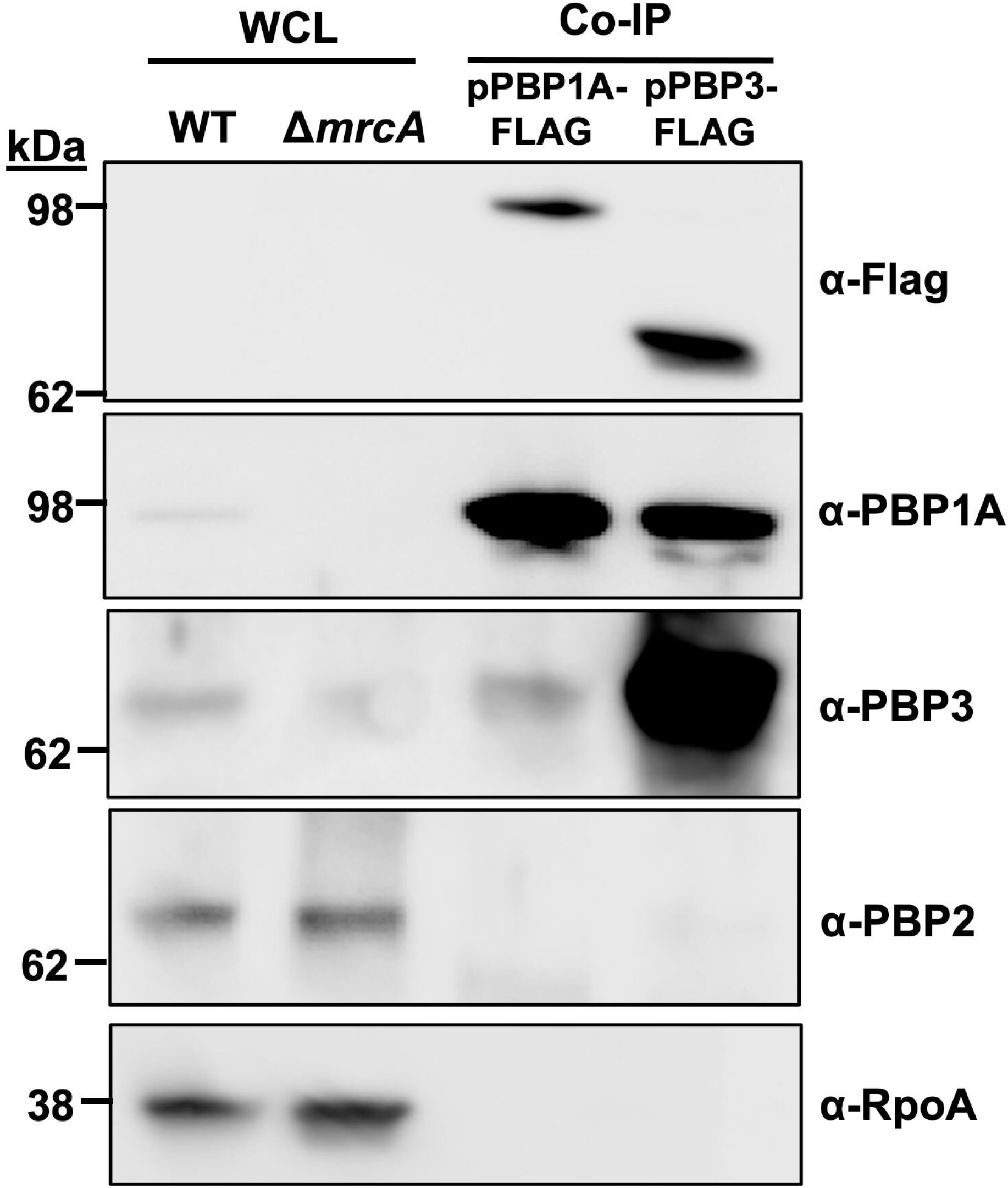
Detection of *in vivo* interactions between PBP1A and PBP3, but not PBP2. Immunoblots with whole-cell lysates (WCL) of wild type (WT) and Δ*mrcA* (lanes 1 and 2) were run as controls. Lanes 3 and 4 contain analysis of PBP1A-FLAG and PBP3-FLAG coimmunoprecipitated (Co-IP) *A. baumannii* proteins. *A. baumannii* cells were cross-linked with DSP (dithiobis(succinimidyl propionate)) to trap complexes and then quenched to stop the cross-linking reaction. Cells were pelleted, osmotically lysed, and then solubilized with a solution containing Triton X-100. After centrifugation, either PBP1A-FLAG or PBP3-FLAG was immunoprecipitated overnight with anti-FLAG M2 affinity gel resin. FLAG-tagged proteins were detected using a monoclonal anti-FLAG antibody. PBP1A, PBP3 and PBP2 were detected with specific antisera after immunoprecipitation. The first two lanes contain proteins from whole cell lysates (WCL) from wild type (WT) or Δ*mrcA A. baumannii*. Antisera specific for RpoA was used as a control to show that cytoplasmic contamination was not present in Co-IP fractions. PBP1A is 94.74 kDa; PBP2 is 74.45 kDa; PBP3 is 67.66 kDa; RpoA is 37.62 kDa.

### PBP3 and PBP5-dependent rescue septation defects in Δ*mrcA A. baumannii*

Prompted by the direct interaction between PBP1A and PBP3, we hypothesized that PBP1A and PBP3 along with its glycosyltransferase partner, FtsW, worked together during septal peptidoglycan biogenesis. We overexpressed PBP3 (**Fig S3B**) or FtsW in Δ*mrcA* cells to determine if either divisome synthase (transpeptidase or glycosyltransferase) could restore the septation defect characteristic of Δ*mrcA* cells (**Fig 4**). Fluorescence and phase microscopy showed that FtsW expression in wild type (WT/pFtsW) produced shorter cells relative to wild type (**Fig 4A**), suggesting the protein was expressed and induced an increased division rate. However, FtsW expression did not rescue the septal defects in Δ*mrcA* cells (Δ*mrcA*/pFtsW) and induced peptidoglycan bulging (**Fig 4B**). In contrast, PBP3 overexpression (Δ*mrcA*/pPBP3_OE_) restored the canonical *A. baumannii* coccobacilli morphology (**Fig 4B**). These data suggest that increased PBP3 activity could compensate for PBP1A defects to promote proper septal peptidoglycan biosynthesis, showing a similar function. Cell length quantifications (**Fig 4C**) also supported our conclusion that PBP3 overexpression rescued septation defects in Δ*mrcA* cells, while FtsW expression did not.

**Figure 4.**
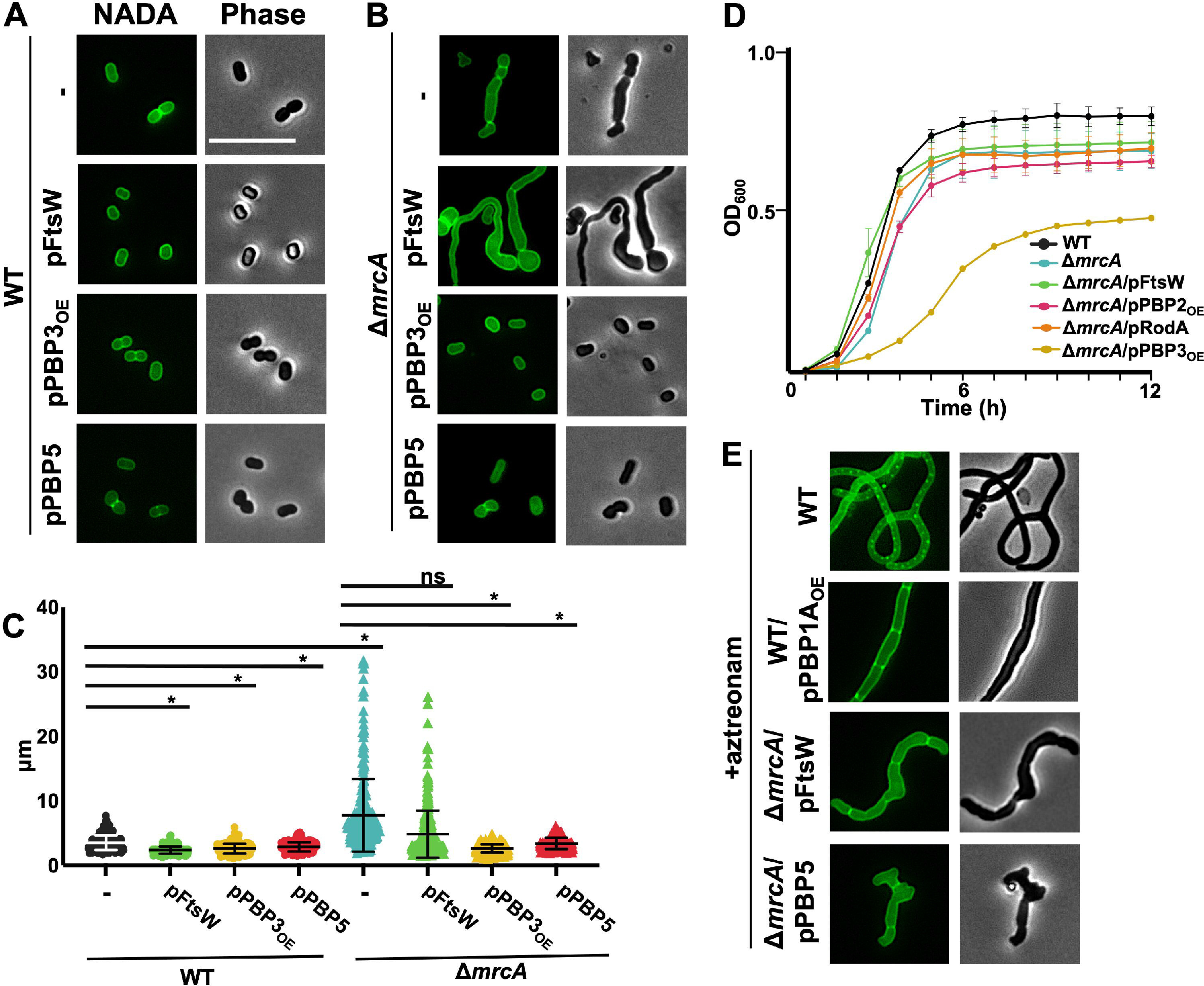
PBP3 and PBP5 rescue division in Δ*mrcA A. baumannii*. (A) Fluorescence and phase microscopy of wild type (WT), WT expressing FtsW, PBP3_OE_, and PBP5. Scale bar = 10 μm. (B) Fluorescence and phase microscopy of Δ*mrcA*, expressing FtsW, PBP3_OE_, and PBP5. (C) Length (pole to pole) (*n* ≥ 300) was calculated using ImageJ software with MicrobeJ plugin. Each dot represents one cell. Significance testing conducted using Student *t* test with two-tailed distribution assuming equal variance. * *P* < 0.05. ns is not significant (D) Optical density (OD_600_) growth curve of wild type (WT), Δ*mrcA*, Δ*mrcA* expressing various peptidoglycan synthases. Error bars indicate standard deviation. (E) Fluorescence and phase microscopy of WT, WT expressing PBP1A_OE_, Δ*pbp1A* expressing FtsW and PBP5, treated with sub-MIC aztreonam (8.0 mg/L).

Furthermore, several studies showed that PBP5, a DD-carboxypeptidase encoded by *dacA*, rescues division defects in PBP3-depleted *E. coli* (1, 36) and other bacteria (37). Considering PBP3 rescued septation defects in Δ*mrcA* cells, we also analyzed if *A. baumannii* PBP5 could also rescue the septation defect in Δ*mrcA* cells. PBP5 expression (Δ*mrcA*/pPBP5) rescued the septation defect in Δ*mrcA* cells, like PBP3 overexpression (**Fig 4B**). Cell length quantifications also supported our conclusion (**Fig 4C**). While only transpeptidase (PBP3 and PBP5) expression rescued the division defect in the Δ*mrcA* strain, both FtsW and transpeptidase overexpression contributed to phenotypic changes in Δ*mrcA A. baumannii*. To determine if these changes were specific to loss of PBP1A activity, we overexpressed FtsW, PBP3, and PBP5 in wild type (**Fig 4A**). Expression of the divisome-associated PG synthases in wild type produced shorter cells (**Fig 4A and 4C**). However, in contrast to PBP1A overexpression which promotes a spherical phenotype (**Fig 2C**), the overexpression of divisome components maintained coccobacilli morphology (**Fig 4A**). Together, these studies imply that in *A. baumannii* PBP1A, PBP3 and PBP5 could work together to coordinate proper septal peptidoglycan biogenesis.

### PBP3 and PBP1A do not share redundant septal peptidoglycan biosynthesis activity

Previous studies from our lab (23) and others (24) demonstrated a significant fitness defect in Δ*mrcA A. baumannii*. We next asked if restoring productive septation in Δ*mrcA* cells would be sufficient to rescue fitness. We overexpressed each of the monofunctional synthases in Δ*mrcA* cells and assessed fitness with optical density growth curves (**Fig 4D**). Intriguingly, PBP3 overexpression (**Fig S3B**) was sufficient to rescue the septation defect in Δ*mrcA* cells (**Fig 4B and 4C**), but not the fitness defect **(Fig 4D)**. However, overexpression of PBP3 in wild type cells also produced a growth defect (**Fig S3C**), suggesting that PBP3 overexpression is toxic regardless of PBP1A activity. Intriguingly, PBP2 expression (**Fig S3D**) did not rescue the Δ*mrcA* fitness defect but was notably toxic when expressed in wild type cells (expressing PBP1A).

We next asked if PBP1A overexpression would similarly rescue division when PBP3 is disrupted. To disrupt PBP3, we treated cells with 0.5-MIC of aztreonam, which has high specificity for PBP3 and effectively prevents division in *A. baumannii* (23) and other Gram-negative bacteria (38), but we also cannot rule out off-target effects. When PBP1A was overexpressed in aztreonam-treated cells, the filamentous phenotype remained **(Fig 4E)**. In contrast to our data showing PBP3 is sufficient to promote division in Δ*mrcA A. baumannii*, PBP1A cannot rescue division when PBP3 is disrupted. Consistent with previous reports in *E. coli* (14, 39), PBP3 is an essential divisome component, where it regulates septal peptidoglycan biogenesis, and its activity cannot be compensated for by PBP1A in *A. baumannii*. Together, these data suggest that while both PBP1A and PBP3 interact in the divisome and possibly have some overlapping functions, they also have distinct roles in septal peptidoglycan biosynthesis. aPBP activity is hypothesized to competitively deplete lipid II precursors (1, 4, 5, 40). Therefore, PBP1A activity may more rapidly direct lipid II to sites of septal peptidoglycan biogenesis relative to FtsW/PBP3.

## Discussion

Peptidoglycan biosynthesis is a validated target to treat Gram-negative infections, and recent studies have highlighted a key role for PBP1A in proper division and fitness in *A. baumannii* (23, 24, 41). These studies strongly imply that the canonical coccobacilli morphology associated with *A. baumannii* is largely dependent on PBP1A peptidoglycan synthase activity at the septum, an unexpected enzymatic role relative to well-studied Gram-negative model systems. In rod-shaped *E. coli*, PBP1A and PBP1B have distinct roles. PBP1A coordinates with elongasome to polymerize peptidoglycan along the side wall (14) and PBP1B interacts with the divisome to assemble septal peptidoglycan (42). However, disruption of either aPBP enzyme is compensated by the other, indicating they are functionally redundant (14). Despite widely accepted dogma in Gram-negative bacteria, PBP1B activity cannot compensate for PBP1A in *A. baumannii*. Intriguingly, the PBP1A regulator, LpoA, an outer membrane lipoprotein that stimulates synthase activity in *E. coli*, is not conserved in *A. baumannii*. Absence of the cognate regulator may support a unique activity for PBP1A, as a septal peptidoglycan synthase. In line with this hypothesis, manipulating PBP1A levels alone are sufficient to induce morphological changes in *A. baumannii* (**Fig 2**). Further, PBP1A is enriched at the midcell during growth (**Fig 2**) where we found it directly interacts with PBP3, but we did not detect a direct PBP2-PBP1A interaction in growth phase (**Fig 3**). Together, these data support a model where PBP1A directly promotes septal peptidoglycan biosynthesis in *A. baumannii*.

We have also shown that PBP1A levels are dynamic, where increased PBP1A levels in stationary phase were evident relative to growth phase (**Fig 1D**), implying it may serve an additional role. Increased PBP1A levels may reflect a final septation event at the end of growth phase before entry into stationary phase. It is also possible that during growth phase PBP1A primarily interacts with the divisome to mediate septation, but increased levels in stationary phase could also promote activity along the cell wall axis in complex with the elongasome or possibly as a repair complex (40, 43). In fact, previous work (44) suggested that PBP1A showed weak interactions with putative elongasome components (not including PBP2). However, there are two concerns with these data, including the two-hybrid screen was not validated and only direct interactions between PBP1A and PBP2 have been described in *E. coli* (17). Consistent with our previous analysis (23) another group also showed (41) that overexpression of PBP1A in wild type cells induced cell rounding (**Fig 2C**). While the other group speculated that PBP1A inhibits elongasome activity (41), it is also possible that PBP1A promotes an increased septation rate to promote formation of short, round cells. It should also be acknowledged that overexpression of PBP1A is not necessarily an indication of increased activity as PBP1A because it traditionally requires and outer membrane lipoprotein activator in other Gram-negative bacteria by the outer membrane lipoprotein LpoA (45). It is unknown how the elongasome and divisome complexes compete for lipid II precursors, but our studies here (**Fig 2A**) suggest that recruitment of PBP1A to the midcell could shift the precursor pool toward septal peptidoglycan synthesis during growth, while indirectly inhibiting lipid II availability to the elongasome. Additional studies are warranted to better understand outcomes of PBP1A activity on elongasome-dependent peptidoglycan biogenesis.

PBP1A overexpression was also not sufficient to compensate for aztreonam-inhibited PBP3 activity. Whereas PBP1A and PBP3 are both necessary for productive septation in wild type *A. baumannii*, these findings indicate the two septal peptidoglycan synthases have independent roles in division. Considering PBP1A localizes to the site of septal peptidoglycan biogenesis prior to septum formation (**Fig 2B**), it may support a model where PBP1A may potentially build the recently described septal peptidoglycan wedge that forms after cellular constriction and prior to septation (4). Previous work suggested that septal peptidoglycan wedge formation prevents lysis during division and may be one possible mechanism for the fitness defect we and others (24) have observed in Δ*mrcA A. baumannii*. Moreover, transpeptidase inactivated PBP1A, but not PBP1B or PBP2, increased cellular lysis in *A. baumannii* under standard growth conditions (24), further supporting PBP1A as a major contributor to septal peptidoglycan biogenesis during division.

Another possible mechanism that PBP1A could contribute to divisome activity could be as an important component of the feedback network in peptidoglycan hydrolase activity. In *E. coli*, several elongasome, divisome and hydrolytic enzymes, including PBP1A, are loosely associated with the outer membrane lipoprotein, NlpI (46). NlpI is thought to act as a scaffold for both hydrolase and synthase complexes (46), and is important for regulating proteolysis of the hydrolytic endopeptidase, MepS (47). NlpI phenotypes in *E. coli* have striking similarities to the *A. baumannii* PBP1A phenotype, where depletion leads to filamentation and overexpression produces rounded, ovoid cells (48). While *A. baumannii* does not encode an NlpI homolog, the phenotypic similarities are intriguing and suggest PBP1A may have an additional, indirect role in localization or regulation of hydrolase activity. Consistent with this hypothesis, Δ*mrcA* forms multiple septal sites, but septation is delayed, suggesting mis-regulation of hydrolase activity.

Not surprisingly, PBP3 overexpression (but not FtsW) rescued the division defect in Δ*mrcA* cells. PBP3 and FtsW are cooperative divisome partners; however, it is not completely unexpected that PBP3 overexpression alone would rescue the division defect. Monofunctional glycosyltransferases require transpeptidases for functionality (11), likely to prevent peptidoglycan polymerization without crosslinking if the transpeptidase is defective or absent. In this context, it is not surprising that overexpression of the transpeptidase, PBP3, was capable of shifting peptidoglycan biogenesis towards division to phenotypically compensate for PBP1A inactivation, while overexpression of FtsW could not. While it is intriguing to speculate that PBP3 and FtsW work together to provide the major septal peptidoglycan polymerization activity and PBP1A works to remodel the synthesized glycan chains, we previously demonstrated that the PBP1A glycosyltransferase activity is necessary for proper septation in *A. baumannii* (23), indicating a direct enzymatic role. Lastly, it is also possible that over expression of FtsW did not increase overall enzymatic activity. In *Bacillus subtilis*, YofA regulates FtsW (49); however, *A. baumannii* lacks a cognate YofA homolog.

PBP5 also rescued the Δ*mrcA* division defects in *A. baumannii*. PBP5 is a DD-carboxypeptidase found in both Gram-negative and Gram-positive bacteria that localizes to peptidoglycan biogenesis sites, where it forms tetrapeptides by cleaving the terminal D-alanine from its pentapeptide substrates (36, 50). While processing pentapeptides to tetrapeptides is important for peptidoglycan maturation, it is not clear why PBP5 overexpression resolves division defects in PBP3-depleted organisms (1, 36, 37), but these studies suggest PBP5 could function similarly in *A. baumannii*. A recent report speculated that following PBP5-dependent tetrapeptide formation, an unidentified periplasmic LD-carboxypeptidase could modify periplasmic tetrapeptides to tripeptides (51), which were proposed to be the primary substrate of PBP3 (1, 52); however, increased PBP3 activity is dependent on lipid II availability, which presumably shifts peptidoglycan biogenesis towards division, away from elongation. If this pathway were intact in *A. baumannii*, it would be possible that PBP5-dependent increases in tripeptide pools could increase substrate availability to PBP3 despite depletion, which could potentially rescue the division defect.

PBP1A expression prevents selection of viable colistin resistant LOS^-^ *A. baumannii* (22). It was recently proposed that PBP1A may directly interfere with the elongasome activity by direct inhibition of PBP2 (41). Supporting this, our previous study (23), demonstrated that PBP1A overexpression promotes cell rounding. However, the more recent study (41) did not provide evidence for interactions between PBP1A and PBP2. In our studies, PBP2 levels in Δ*mrcA* were not increased relative to wild type. However, a curious finding suggested that PBP2 overexpression in PBP1A-producing *A. baumannii* led to rapid lysis (data not shown). In contrast, PBP2 overexpression in Δ*mrcA* was tolerated (**Fig S3C**). In Δ*mrcA A. baumannii*, PBP2 overexpression toxicity may be alleviated because competition from the divisome (lacking PBP1A) for lipid II substrate is reduced and septation is slowed. While these curious findings need to be explored further, preliminary data support a model where increased lipid II availability to PBP2 may result from defects in PBP1A activity. We plan to explore how this peptidoglycan regulatory mechanism promotes *A. baumannii* survival when the outer membrane is severely defective because it could provide novel insights into antibiotic resistance mechanisms.

## Materials and Methods

### Bacterial Strains and Growth

All primers are listed in Table S1, and strains and plasmids are listed in Table S2. All *A. baumannii* strains were grown from freezer stocks initially on Luria-Bertani (LB) agar at 37° C. For selection, 25 μg/ml of kanamycin was used when appropriate. Strains that harbored the pMMB plasmid for coIP, complementation or overexpression were supplemented with 25 μg/ml of kanamycin and 2 mM isopropylthio-β-galactoside (IPTG).

### Fluorescent NADA staining

As previously described (23, 27), overnight cultures were back-diluted to OD_600_ 0.05 and grown at 37°C in Luria broth until they reached stationary or mid-logarithmic growth phase. Cells were washed once with Luria broth and normalized to OD_600_ 1.0. 3 μl of 10 mM of NBD-(linezolid-7-nitrobenz-2-oxa-1,3-diazol-4-yl)-amino-D-alanine (NADA) (Tocris Bioscience) was added to the resuspension. Cells were incubated with NADA at 37° C for 0.5 hours. Following incubation, cells were washed once and fixed with 1x phosphate buffered saline containing a (1:10) solution of 16 % paraformaldehyde.

### Microscopy

Fixed cells were immobilized on agarose pads and imaged using an inverted Nikon Eclipse Ti-2 widefield epifluorescence microscope equipped with a Photometrics Prime 95B camera and a Plan Apo 100 m X 1.45 numerical aperture lens objective, as previously described (23, 27). Green fluorescence and red fluorescence images were taken using a filter cube with a 470/40 nm or 560/40 nm excitation filters and 632/60 or 535/50 emission filters, respectively. Images were captured using NIS Elements software.

### Image analysis

All images were processed and pseudo-colored with ImageJ FIJI (53) and MicrobeJ plugin was used for quantifications of pole-to-pole lengths (54). For pole-to-septa length quantifications, a segmentation code written for Image J was used (see Appendix 1). Cell length, width and fluorescence data were plotted in Prism 9 (GraphPad 9.3.1). Demographs were generated using MicrobeJ plugin. 100 cells were analyzed for pole-to-septa lengths, 450 cells were used for demographs and ≥300 cells were analyzed for all other experiments. Each experiment was independently replicated three times, and each replication was used in the dataset. One representative image from the datasets were included in the figure.

### Growth curves

Growth curves were performed as previously described (23, 55, 56). Briefly, overnight cultures were back diluted to OD_600_ 0.01 and set up as triplicate biological replicas in either a 96-well plate or 24-well plate (BrandTech BRAND). A BioTek SynergyNeo^2^ microplate reader was used to record optical density, which was read at OD_600_ every hour. The microplate reader was set to 37°C with continuous shaking. Growth curves were plotted in Prism 9. Each growth curve experiment was independently replicated three times and one representative dataset was reported.

### Western blotting

Western blot analysis was carried out via gel transfer to polyvinylidene fluoride (PVDF) (Thermo-Fisher Scientific). All blots were blocked in 5% milk for 2 hours. The primary antibodies α-PBP1A, α-PBP3, α-PBP2, and α-RpoA were used at 1:1000, 1:500, 1:300, and 1:1000, respectively followed by α-rabbit-HRP secondary antibody at 1:10,000 (Thermo-Fisher Scientific). Supersignal West Pico PLUS (Thermo-Fisher Scientific) was used to measure relative protein concentrations.

### Co-immunoprecipitation

The protocol was adapted from previous work (33). Briefly, cultures were initially grown on LB agar with overnight at 37° C. A single colony was used to inoculate 5 ml Luria broth and grown overnight at 37° C. The overnight broth was diluted back to OD_600_ 0.05 and grown to mid-logarithmic growth phase at 37° C in 50 ml Luria broth. Cultures were collected, washed with 1x PBS, and incubated with 25 mM DSP (Thermofisher) in a total volume of 1 ml PBS for 1 hour with shaking at 37° C. The reaction was quenched with room temperature incubation of 400 μl 1 M glycine with rocking for 15 minutes. Cells were lysed and solubilized overnight at 4° C with rocking. Lysates supernatant was centrifuged twice at 10,000 X g for 20 minutes and the pellet was discarded. 30 μl of ANTI-FLAG® M2 Affinity Resin (Sigma) was added to the supernatant and incubated overnight at 4° C with rocking. Resin was harvested at 6,000 X g and washed 4 times with RIPA buffer. Pellet resuspended in 100 μl 1x SDS-Page loading buffer with 5% BME. Samples were boiled for 7 minutes and used for western blotting.

## Acknowledgements

This work was supported by funding from the National institutes of Health (grants AI168159 and GM143053 to JMB)

## Notes

### Competing Interest Statement

The authors have declared no competing interest.

