## Supplemental material for "PBP1A directly interacts with the divisome complex to promote septal peptidoglycan synthesis in *Acinetobacter baumannii*"

**Table S1:** Primers used in this study

| **Primer** | Sequence 5' -> 3' | **Description** | **Used to make** |
|---|---|---|---|
| RodA Forward | CGCGAGCTCATGTCTCCTAGTCCAC A | [CGC]-[SacI]-[RodA] | pRodA |
| RodA Reverse | CGCGTCGACTTATCGATGTGTATGA ATA | [CGC]-[SalI]-[RodA] | pRodA |
| FtsW Forward | CGCGGTACCATGGCAGGCTTA | [CGC]-[KpnI]-[Ftsw] | pFtsW |
| FtsW Reverse | CGCGTCGACTTAGAAGTTTGATTCT T | [CGC]-[Sal]-[Ftsw] | pFtsW |
| PBP2 Forward | CGCGGTACCATGAAACAGCACTTTC CT | [CGC]-[KpnI]-[PBP2] | pPBP2$_{OE}$ |
| PBP2 Reverse | CGCGGATCCTTATTTATCATCATCA TCTTTATAATCTTCATCGACCTCGTT | [CGC]-[BamHI]-[FLAG]-[PBP2] | pPBP2$_{OE}$ |
| PBP3 Forward | CGCGGTACCATGGTAGATAAGCGA ACAAAGCAAACACG | [CGC]-[KpnI]-[PBP3] | pPBP3-FLAG |
| PBP3 Reverse | CGCGTCGACTTACTTGTCGTCATCG TCTTTGTAGTCCCTGCGAATAGGAT | [CGC]-[SalI]-[FLAG]-[PBP3] | pPBP3-FLAG |
| PBP1A-FLAG Forward | CGCCTCGAGATGAAAAAGCTATCC AGTTTGGGCTT | [CGC]-[XhoI]-[PBP1A] | pPBP1A-FLAG |
| PBP1A-FLAG Reverse | CGCGGTACCTTATTTATCATCATCA TCTTTATAATCCTCAATTTGATTAA T | [CGC]-[KpnI]-[FLAG]-[PBP1A] | pPBP1A-FLAG |
| pMMB sequencing Forward | CGGTTCTGGCAAATATTCTGAAA | Plasmid confirmation primer | pRodA, pFtsW, pPBP2-FLAG, pPBP3-FLAG, pPBP1A-FLAG |
| pMMB sequencing Reverse | GCCGCCAGGCAAATTCTGTT | Plasmid confirmation primer | RodA, pRodA, pFtsW, pPBP2-FLAG, pPBP3-FLAG, pPBP1A-FLAG |
| PBP2 FRT 5' | ACAAAGTCAAAAAAGCCAATTTAC CTCTTCACTGAAACTTTGAAAAATA TGCGATATTTTAAGTGCGCTTGCGT TAGAATAAGCAGCTATTTTTCTCAC | 5' Recombineering primer | Δ*mrdA* |

| | | | |
|---|---|---|---|
| | CCTATGTCATTAAGTCGATCCGTATG | | |
| PBP2 FRT 3’ | CATGAGCCAGTTAAATAAAGTTTCGCGGACACGATCGGGAGTTGGTCTTAATCCTTCAATACTGGCGAATGGTAAAACTCGTCTTTTCCATTCGCCCCCAATAATGCGTAATTGATTTTTCATTA | 3’ Recombineering primer | Δ*mrdA* |
| PBP2 Forward | GCTTACTGTCAAAACTGCAAGTAAGGG | Mutation confirmation primer | Δ*mrdA* |
| PBP2 Reverse | TGTTCTTTTAGGCGTGGTAAGGCT | Mutation confirmation primer | Δ*mrdA* |

**Table S2:** Strains and plasmids used in this study

| Strain/Plasmid | Description | Reference/Source |
|---|---|---|
| **Strains** | | |
| *A. baumannii* ATCC 17978 | wild type | ATCC (1) |
| *A. baumannii* ATCC 17978 | Δ*mrcA* | (2) |
| *A. baumannii* ATCC 17978 | Δ*mrcA*/pPBP1A | (2) |
| *A. baumannii* ATCC 17978 | WT/pPBP1A$_{OE}$ | (3) |
| *A. baumannii* ATCC 17978 | Δ*mrcA*/pPBP1A$_{OE}$ | (3) |
| *A. baumannii* ATCC 17978 | Δ*mrcB* | (2) |
| *A. baumannii* ATCC 17978 | Δ*mrcA*/pPBP1A-mCherry | (3) |
| *A. baumannii* ATCC 17978 | Δ*mrcA*/pPBP1B$_{OE}$ | This Study |
| *A. baumannii* ATCC 17978 | Δ*mrcA*/pFtsW | This Study |
| *A. baumannii* ATCC 17978 | Δ*mrcA*/pPBP2$_{OE}$ | This Study |
| *A. baumannii* ATCC 17978 | Δ*mrcA*/pPBP5 | This Study |
| *A. baumannii* ATCC 17978 | Δ*mrcA*/pPBP3$_{OE}$ | This Study |

| | | |
|---|---|---|
| *A. baumannii* ATCC 17978 | WT/pPBP1A-FLAG | This Study |
| *A. baumannii* ATCC 17978 | WT/pPBP3-FLAG | This Study |
| *A. baumannii* ATCC 17978 | WT/pPBP3$_{OE}$ | This Study |
| *A. baumannii* ATCC 17978 | WT/pFtsW | This Study |
| *A. baumannii* ATCC 17978 | WT/pPBP5 | This Study |
| *A. baumannii* ATCC 17978 | Δ*mrdA* | This Study |
| **Plasmids** | | |
| pABBRKn | pABBR_ MCS with the $Kn^R$ gene from pKD4 replacing the *bla* gene, $Kn^R$ | (2) |
| pMMBKn | pMMBKn_MCS with the $Kn^R$ gene from pKD4 replacing the *bla* gene, $Kn^R$ | (2) |
| pABBRKn-mCherry | pABBRKn with the *mCherry gene* inserted into the KpnI and SacI sites, $Kn^R$ | (3) |
| pPBP1A | pABBRKn with the *mrcA* gene and native promoter inserted into the XhoI and KpnI sites, $Kn^R$ | (3) |
| pPBP1A$_{OE}$ | pMMBKn with the *mrcA* gene inserted into the XhoI and KpnI sites, under an IPTG inducible promoter, $Kn^R$ | (3) |
| pPBP1A-mCherry | pABBRKn-mCherry with the *mrcA* gene inserted into the XhoI and SacI sites, $Kn^R$ | (3) |
| pAT03 | pMMB67EH with FLP recombinase, $Amp^R$ | (4) |
| pAT04 | pMMB67EH with the REC$_{Ab}$ system, $Tet^R$ | (4) |
| pKD4 | $Kn^R$ | (4) |
| pPBP1B$_{OE}$ | pMMBKn with the *mrcB* gene inserted into the KpnI and SalI sites, $Kn^R$ | (2) |
| pFtsW | pMMBKn with the *ftsW* gene inserted into the KpnI and SalI sites, $Kn^R$ | This Study |
| pPBP5 | pMMBKn with the *dacA* gene inserted into the KpnI and SalI sites, $Kn^R$ | This Study |
| pPBP2$_{OE}$ | pMMBKn with the *mrdA* gene with a FLAG tag inserted into the KpnI and BamHI sites, $Kn^R$ | This Study |
| pPBP3$_{OE}$ | pMMBKn with the *ftsI* gene with a FLAG tag inserted into the KpnI and BamHI sites, $Kn^R$ | This Study |

| | | |
|---|---|---|
| pPBP3-FLAG | pMMBKn with the *ftsI* gene with a FLAG tag inserted into the KpnI and SalI sites, $Kn^R$ | This Study |
| pPBP1A-FLAG | pMMBKn with the *mrcA* gene with a FLAG tag inserted into the XhoI and KpnI sites, under an IPTG inducible promoter, $Kn^R$ | This Study |

**Supplemental Figures**

**Figure S1**

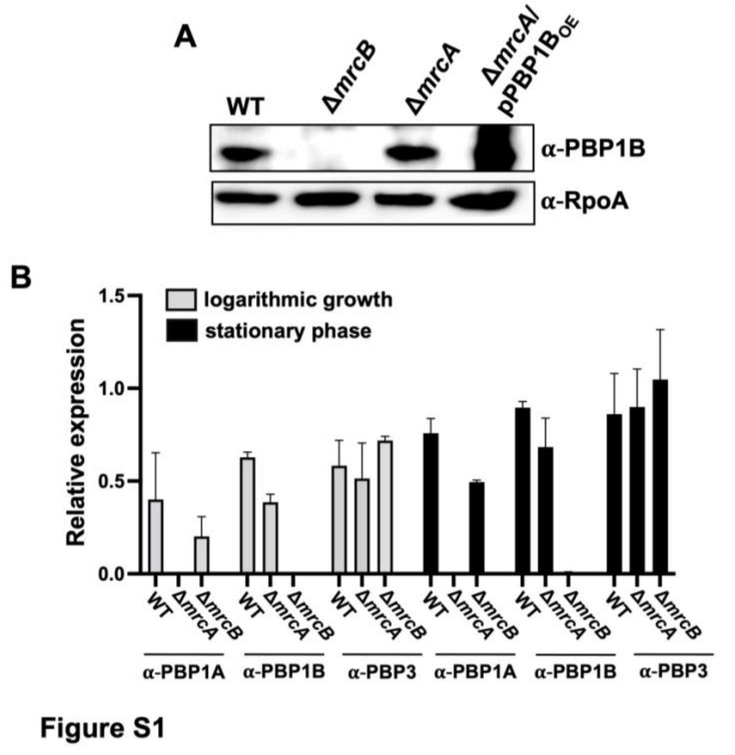


**Figure S1:** (A) PBP1B expression levels in *A. baumannii* wild type (WT) and mutants. RpoA is the loading control. PBP1B is 88.21 kDa; RpoA is 37.62 kDa. (B) Densitometry of western blot analysis as calculated by ImageJ. Relative protein levels are represented as fold change over RpoA loading control. Error bars indicate the variance between two independent experiments. A representative image is shown in Fig 1D.

**Figure S2**

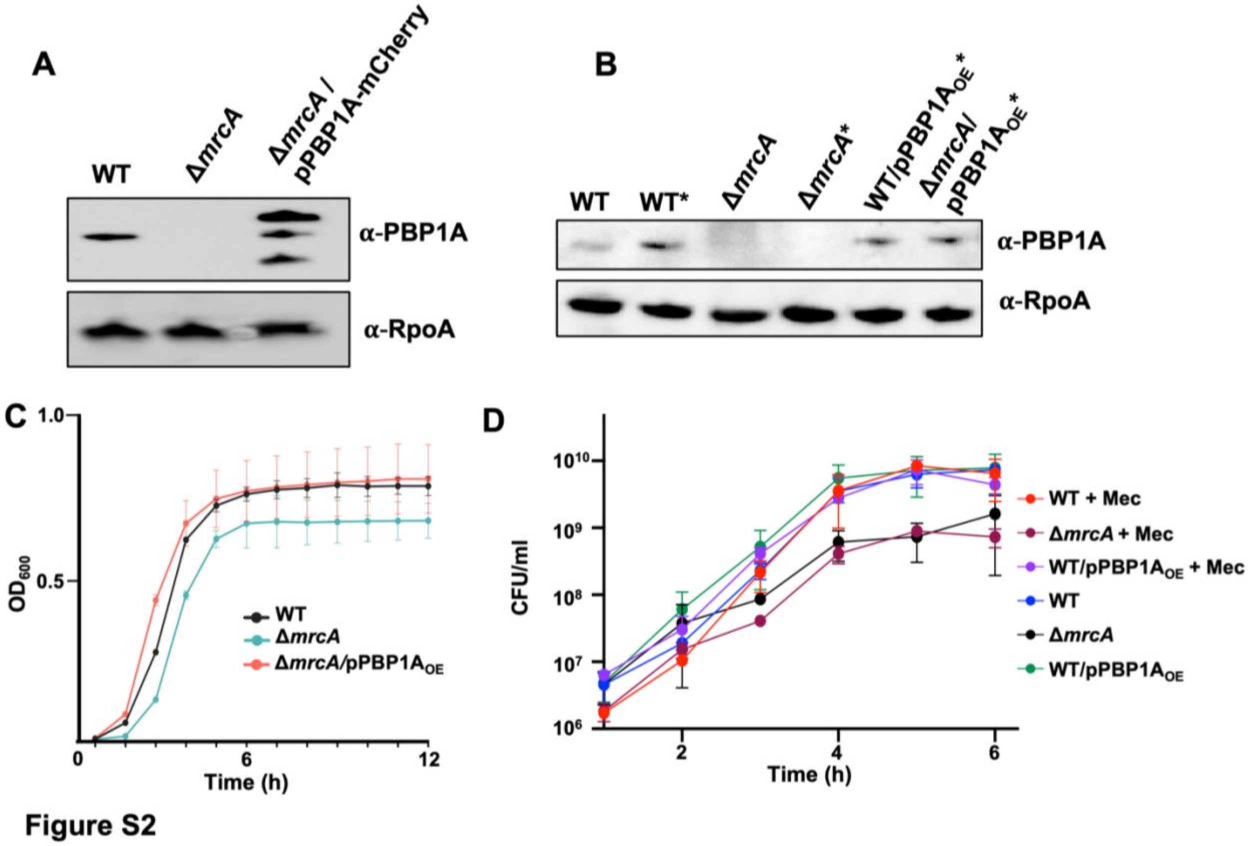


**Figure S2:** (A) PBP1A-mCherry expression under native promoter in wild type (WT) *A. baumannii*. The top band corresponds with the PBP1A-mCherry fusion predicted molecular weight. RpoA is the loading control. PBP1A is 94.74 kDa; PBP1A-mCherry is 120.01 kDa; RpoA is 37.62 kDa. (B) PBP1A expression in *A. baumannii* WT and mutants. * = treated with sub-MIC mecillinam (32 mg/L). RpoA is the loading control. (C) PBP1A overexpression restores the *mrcA* growth defect. Optical density growth curve of WT, Δ*mrcA*, and Δ*mrcA*/pPBP1A$_{OE}$ showing that PBP1A expression restores the growth (fitness) defect. Error bars represent standard deviation. (D) CFU/ml of WT, Δ*mrcA*, and WT/pPBP1A$_{OE}$ grown in the presence or absence of sub-MIC mecillinam (32 mg/L). Error bars represent standard deviation.

**Figure S3**

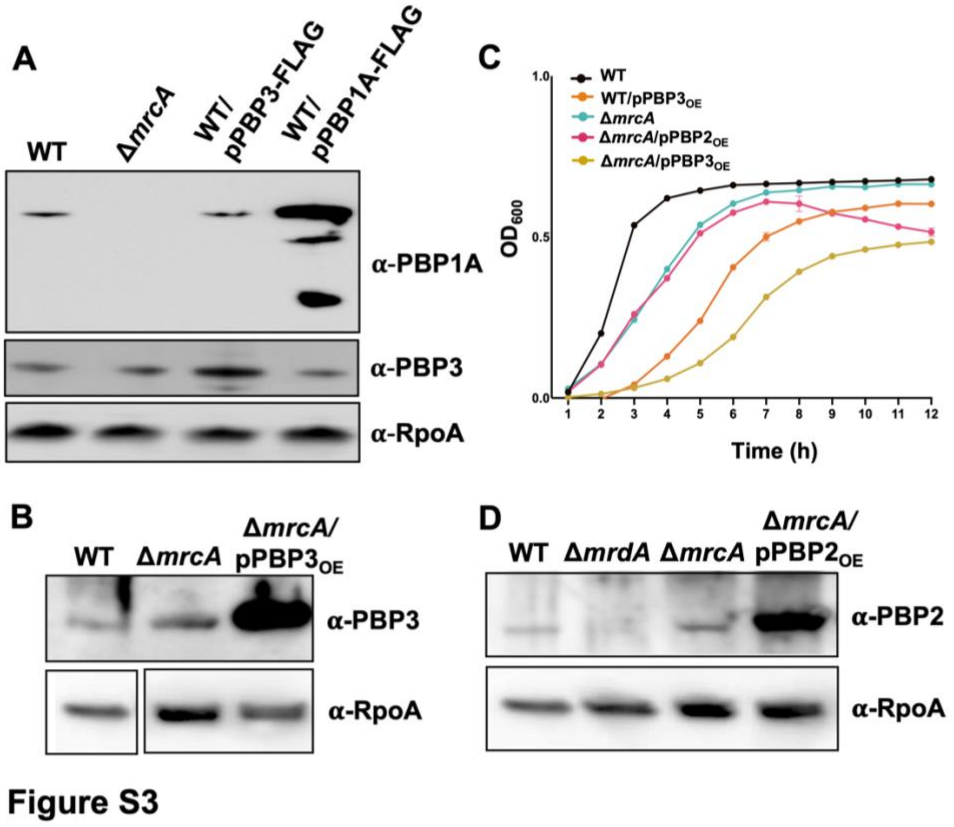


**Figure S3:** (A) Whole-cell lysates of FLAG fusion strains included in the co-immunoprecipitation to determine relative protein abundance. PBP1A is 94.74 kDa; PBP3 is 67.66 kDa; RpoA is 37.62 kDa. (B) PBP3 expression in *A. baumannii* wild type (WT) and mutants. (C) Growth curve of wild type and Δ*mrcA* overexpressing PBP3 and PBP2. Error bars represent standard deviation. (D) PBP2 expression in *A. baumannii* WT and mutants. PBP2 is 74.45 kDa.
